# ZIF-1-mediated degradation of endogenous and heterologous zinc finger proteins in the *C. elegans* germ line

**DOI:** 10.1101/2023.07.10.548405

**Authors:** Aaron Z.A. Schwartz, Yusuff Abdu, Jeremy Nance

## Abstract

Rapid and conditional protein depletion is the gold standard genetic tool for deciphering the molecular basis of developmental processes. Previously, we showed that by conditionally expressing the E3 ligase substrate adaptor ZIF-1 in *Caenorhabditis elegans* somatic cells, proteins tagged with the first CCCH Zn finger (ZF1) domain from the germline regulator PIE-1 degrade rapidly, resulting in loss-of-function phenotypes. The described role of ZIF-1 is to clear PIE-1 and several other CCCH Zn finger proteins from early somatic cells, helping to enrich them in germline precursor cells. Here, we show that proteins tagged with the PIE-1 ZF1 domain are subsequently cleared from primordial germ cells in embryos and from undifferentiated germ cells in larvae and adults by ZIF-1. We harness germline ZIF-1 activity to degrade a ZF1-tagged heterologous protein from PGCs and show that its depletion produces phenotypes equivalent to those of a null mutation. Our findings reveal that ZIF-1 switches roles from degrading CCCH Zn finger proteins in somatic cells to clearing them from undifferentiated germ cells, and that ZIF-1 activity can be harnessed as a new genetic tool to study the early germ line.

## Introduction

The ability to silence gene products is critical for investigating cellular and developmental processes. Commonly employed silencing methods include conditionally mutating endogenous genes or depleting their mRNA products using RNA interference. However, phenotypes do not arise when using these approaches until any existing protein gene product decays, limiting their utility in studying rapidly changing developmental events (Nance and Frokjaer-Jensen 2019). Several methods have been developed to directly deplete protein gene products. The auxin-inducible degron system, which has been used in a wide variety of model systems from yeast to mammals (Nishimura *et al*. 2009; Holland *et al*. 2012), is perhaps the most widely employed targeted protein degradation method. In this system, an auxin-inducible degron (AID) domain is fused to a protein of interest. Co-expressing the *Arabidopsis* E3 ligase substrate adaptor TIR1 recruits the AID-tagged protein to an SCF (SKP1–CUL1–F-box) E3 ubiquitin ligase complex when the phytohormone auxin is present, resulting in its ubiquitylation and proteasome-mediated degradation (Nishimura *et al*. 2009; Holland *et al*. 2012). The AID system has been widely used and optimized in *C. elegans* to study gene function in larvae and adults (Zhang *et al*. 2015; Martinez *et al*. 2020; Ashley *et al*. 2021; Hills-Muckey *et al*. 2022; Negishi *et al*. 2022; Sepers *et al*. 2022; Xiao *et al*. 2023). However, auxin appears to have difficulty crossing the embryonic eggshell, and AID-tagged protein knockdown in embryos occurs with variable success (Zhang *et al*. 2015; Negishi *et al*. 2019; Negishi *et al*. 2022). Although custom-modified auxin analogs with increased cell permeability improve degradation in embryos, (Negishi *et al*. 2019; Negishi *et al*. 2022) the AID system is not yet widely used to study embryonic development.

Previously, we developed a degron method called the ZIF-1/ZF1 degron system by repurposing an endogenous *C. elegans* protein degradation pathway (Armenti *et al*. 2014). ZIF-1 is a SOCS-box E3 ligase substrate adaptor protein that helps separate somatic and germline cell fates during early embryogenesis (DeRenzo *et al*. 2003). The germ line arises as the result of four asymmetric cell divisions, beginning with cleavage of the zygote; each division produces a germline precursor cell (P1 – P4) and a somatic sister cell. P4 then divides symmetrically to produce two primordial germ cells (PGCs, named Z2 and Z3) (Sulston *et al*. 1983), which remain quiescent for the remainder of embryogenesis (Fukuyama *et al*. 2006). Several maternally inherited proteins that contain CCCH Zn finger domains become enriched in germline precursor cells, including MEX-1, POS-1, and the transcriptional inhibitor PIE-1 (Mello *et al*. 1996; Seydoux 1996; Guedes and Priess 1997; Tabara *et al*. 1999). ZIF-1 binds to the Zn finger 1 (ZF1) domain of MEX-1, POS-1, and PIE-1 to deplete these proteins from early embryonic somatic cells (Reese *et al*. 2000; DeRenzo *et al*. 2003). Although endogenous ZIF-1 activity disappears from somatic lineages after early embryogenesis (Nance *et al*. 2003), conditionally expressing ZIF-1 in later embryos causes proteins tagged with the PIE-1 ZF1 domain to degrade rapidly, resulting in loss-of-function phenotypes (Armenti *et al*. 2014). Because the ZIF-1/ZF1 degron system can clear a targeted protein quickly, it has been used primarily to study the functions of essential genes in defined embryonic developmental events (Nance and Frokjaer-Jensen 2019).

In contrast to early embryonic somatic cells, ZIF-1 is either absent or inactive in P1 – P4, ensuring that MEX-1, POS-1, and PIE-1 are able to enrich in germline precursor cells (Reese *et al*. 2000; DeRenzo *et al*. 2003) (FIG1A). Shortly after the PGCs are born, however, PIE-1 disappears from these cells (Mello *et al*. 1996; Seydoux and Dunn 1997; Schaner *et al*. 2003) in a *zif-1*-dependent manner (Checchi and Kelly 2006). Thus, ZIF-1 either has an indirect role in PIE-1 stability in PGCs, or ZIF-1 becomes expressed or activated in PGCs and directly targets PIE-1 for degradation.

To better understand when and where proteins tagged with the ZF1 degron are targeted by endogenous ZIF-1, we examined ZIF-1 function in the germline lineage. We show that ZIF-1 is expressed zygotically in embryonic PGCs as well as in undifferentiated germ cells within larvae and adults, where it functions to clear ZF1-tagged proteins from these cells. By harnessing germline ZIF-1 activity, we demonstrate that endogenously tagged proteins fused to the ZF1 domain can be degraded in PGCs to produce loss-of-function phenotypes. Our findings provide new insights into how CCCH Zinc finger proteins are temporally and spatially regulated within the germ line, and introduce a novel genetic tool to investigate the biology of *C. elegans* PGCs and undifferentiated germ cells.

## Results

### ZIF-1 degrades ZF1-tagged proteins in PGCs

To learn if ZIF-1 directly clears proteins containing CCCH Zn finger domains from PGCs, we tested whether fusing the PIE-1 ZF1 domain to a heterologous protein would force the fusion protein to degrade. Rho GTPases CDC-42 (Zilberman *et al*. 2017) and CED-10 (this study; see Methods) endogenously tagged with the ZF1 domain and YFP are expressed ubiquitously in both the soma and germ line. In early embryos, ZF1-YFP-CDC-42 was degraded from somatic cells but persisted in newly born PGCs (**FIG1B**). Like PIE, ZF1-YFP-CDC-42 and ZF1-YFP-CED-10 disappeared from slightly older PGCs of bean stage embryos (**FIG1C,H).** Because ZIF-1 is either absent or inactive in the soma of older embryos (Nance *et al*. 2003; Armenti *et al*. 2014), ZF1-YFP-CDC-42 and ZF1-YFP-CED-10 reappeared in somatic cells of larvae as a result of their zygotic expression **(FIG1D,I)**, but both proteins remained absent from PGCs through the L1 larval stage **(FIG1D,I)**. Transgenic mCherry-PH-ZF1 (Beer *et al*. 2018) expressed from the germline-specific *pie-1* promoter showed a similar pattern of degradation in early embryos and PGCs, except that mCherry-PH-ZF1 did not reappear in the soma **(FIG1L-M**). In *zif-1* mutant embryos, ZF1-YFP-CDC-42, ZF1-YFP-CED-10, and mCherry-PH-ZF1 were present in all embryonic cells, including PGCs (**FIG1E-G, J-K, N-Q)**. Taken together, these findings show that proteins containing a ZF1 domain are degraded in early embryonic somatic cells, protected in germline precursor cells and newly born PGCs, and subsequently degraded by ZIF-1 in slightly older PGCs.

**Figure 1.**
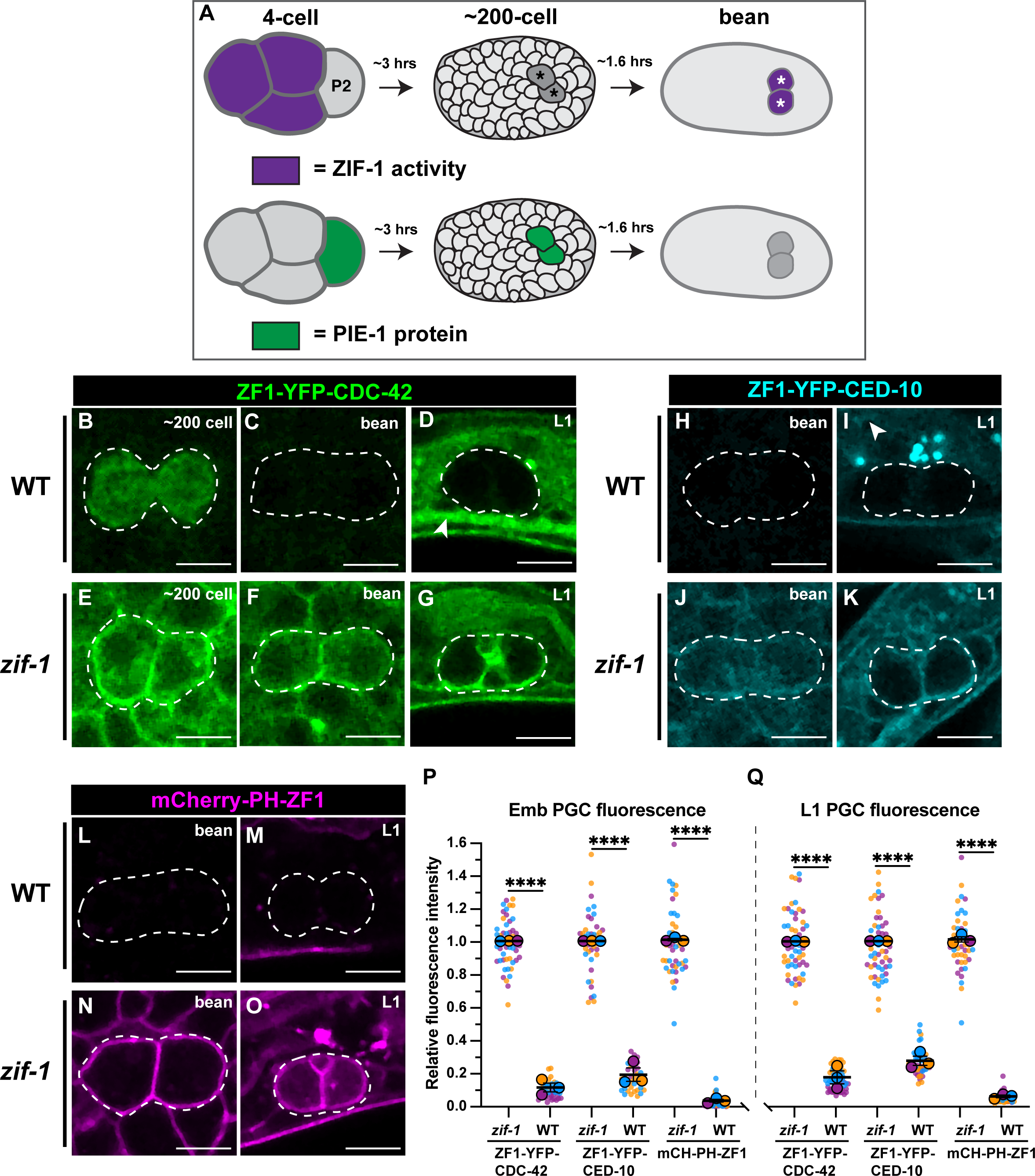
Endogenous ZIF-1 degrades heterologous ZF1-tagged proteins in PGCs. (**A**) Schematic of ZIF-1 activity (purple) and PIE-1 protein (green) localization during early embryogenesis; P2 indicates a germline precursor cell, asterisks mark PGCs. (**B-D**) Localization pattern of ZF1-YFP-CDC-42 in wild-type PGCs (dashed outlines in this and subsequent panels); arrowhead indicates somatic re-expression of ZF1-YFP-CDC-42. (**E-G**) Localization pattern of ZF1-YFP-CDC-42 in *zif-1* mutants. (**H-I**) Localization pattern of ZF1-YFP-CED-10 in wild-type PGCs; arrowhead indicates somatic re-expression of ZF1-YFP-CED-10 (bright spots in (I) are gut granules). (**J-K**) Localization pattern of ZF1-YFP-CED-10 in *zif-1* mutant PGCs. (**L-M**) Localization pattern of mCherry-PH-ZF1 in wild-type PGCs. (**N-O**) Localization pattern of mCherry-PH-ZF1 in *zif-1* mutant PGCs. (**P-Q**) Quantification of fluorescence in bean stage (P) and L1 PGCs (Q) in wild-type and *zif-1* mutants, normalized to expression in *zif-1* mutants. Individual data points from three independent color-coded biological replicates are shown as small dots, the mean from each experiment shown as a larger circle, the mean of means as a horizonal line, and the S.E.M as error bars. \*\*\*\**p* ≤ 0.0001, unpaired two-tailed Student’s *t*-test. Scale bar, 5µm.

### zif-1 is expressed zygotically in PGCs

We next investigated how *zif-1* becomes active in PGCs. 3’ UTR elements regulate the temporal and spatial expression of many germline genes (Merritt *et al*. 2008). A previously characterized GFP-tagged histone H2B reporter controlled by the *pie-1* germline promoter and *zif-1* 3’ UTR (*pie-1*p::*GFP-H2B*::*zif-1^3’UTR^*) is detected in early embryonic somatic cells but not in germline precursor cells (Guven-Ozkan *et al*. 2010). The expression of this reporter has not been described in PGCs. We observed low and variable expression of *pie-1*p::*GFP-H2B*::*zif-1^3’UTR^* in newly born PGCs (5/10 embryos), but not in PGCs from later embryos (0/15 at bean stage) or L1 larvae (0/16) (**FIG2A-C**). Together with previously published findings, these observations indicate that the *zif-1* 3’ UTR can support translation of mRNAs in early somatic cells and in PGCs but not in germline precursor cells, consistent with the pattern of ZIF-1-mediated degradation that we observe. However, they do not address whether the normal source of *zif-1* mRNAs in PGCs results from maternal and/or zygotic transcription.

**Figure 2.**
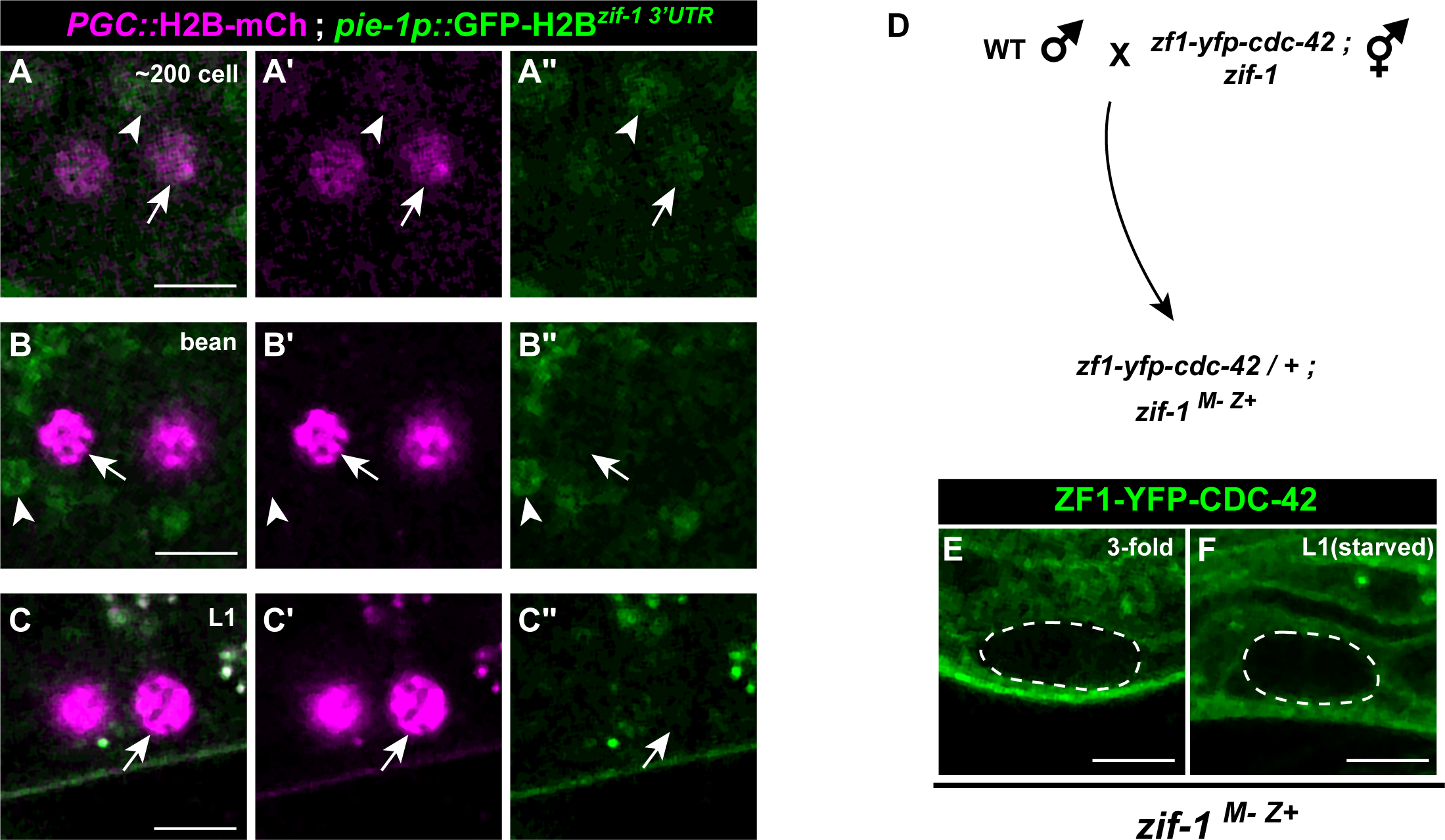
Control of *zif-1* expression. (**A-A’’**) Expression pattern of *pie-1p::GFP-H2B^zif-1^ ^3’UTR^* in early PGCs. PGCs marked with *PGC*::*mCherry-H2B* (*mex5p::mCherry-H2B^nos-2^ ^3’UTR^*). *pie-1p::GFP-H2B^zif-1^ ^3’UTR^* translation is faintly detectable in somatic cells (arrowhead) and PGCs (arrow) in 5/10 embryos. **(B-C’’)** In bean stage embryo and L1 PGCs, *pie-1p::GFP-H2B^zif-1^ ^3’UTR^* is undetectable (arrow); somatic *pie-1p::GFP-H2B^zif-1^ ^3’UTR^* expression is visible in bean stage embryo somatic cells (arrowhead in C-C’’) but not in L1 larvae. (**D**) Crossing scheme to introduce functional *zif-1(+)* zygotically in PGCs. (**E-F**) ZF1-YFP-CDC-42 is lost in embryonic and L1 larval PGCs (dashed outline) when *zif-1* is introduced zygotically (compare to Fig. 1F,G).

We next asked if zygotic *zif-1* transcription is triggered in PGCs. Initially, we considered this possibility unlikely, as only a few genes have been conclusively shown to be zygotically transcribed in PGCs (Subramaniam and Seydoux 1999; Kawasaki *et al*. 2004; Mainpal *et al*. 2015). In addition, ZIF-1 itself is required to degrade PIE-1 (Checchi and Kelly 2006), which blocks transcription by inhibiting RNA Pol II phosphorylation (Seydoux 1996; Seydoux and Dunn 1997). To determine whether *zif-1* is transcribed zygotically in PGCs, we crossed *zif-1* mutant mothers containing the *zf1-yfp-cdc-42* allele with wild-type [*zif-1(+)*] males, and examined whether ZF1-YFP-CDC-42 degraded in PGCs of the F1 *zif-1(M-Z+)* embryos; from this crossing scheme, functional ZIF-1 in F1 embryos could only arise if it were expressed zygotically from the *zif-1(+)* allele introduced by the male (**FIG2D**). Similar to ZF1-YFP-CDC-42 in a wild-type background, but in contrast to ZF1-YFP-CDC-42 in a *zif-1* mutant background, ZF1-YFP-CDC-42 was degraded from late-embryonic (10/10) and L1 larval (7/7) PGCs in *zif-1(M-Z+)* animals (**FIG2E**). Taken together, these findings indicate that *zif-1* is transcribed zygotically in embryonic PGCs, causing proteins containing the ZF1 domain to degrade.

### ZIF-1 remains active in undifferentiated germ cells in larvae and adults

PIE-1 and several other ZIF-1 targets containing CCCH Zinc finger domains are present in portions of the adult germ line (Guedes and Priess 1997; Tenlen *et al*. 2008; Tsukamoto *et al*. 2017; Kim *et al*. 2021), indicating that ZIF-1 germline activity or expression must be inhibited before these proteins are expressed. To determine when after PGC formation ZIF-1 becomes inactive, we examined ZF1-YFP-CDC-42 expression in the larval germ line. Germ line expansion commences when L1 larvae encounter food, which triggers PGCs to re-enter the cell cycle and transition into proliferating germline stem cells (GSCs) (**FIG3A**) (Fukuyama *et al*. 2006; Hubbard and Schedl 2019). In later larvae and adults, GSCs and their proliferating progeny (collectively referred to as ‘undifferentiated germ cells’ hereafter) reside in the distal portion of each arm of the bi-lobed gonad (Hubbard and Schedl 2019) (**FIG S1A-I**); undifferentiated germ cells move more proximally within the gonad as a result of proliferation and eventually differentiate by entering meiosis. The first meiotic germ cells appear in L3-stage larvae (Kimble and White 1981; Pepper *et al*. 2003; Hansen *et al*. 2004; Hubbard and Schedl 2019).

**Figure 3.**
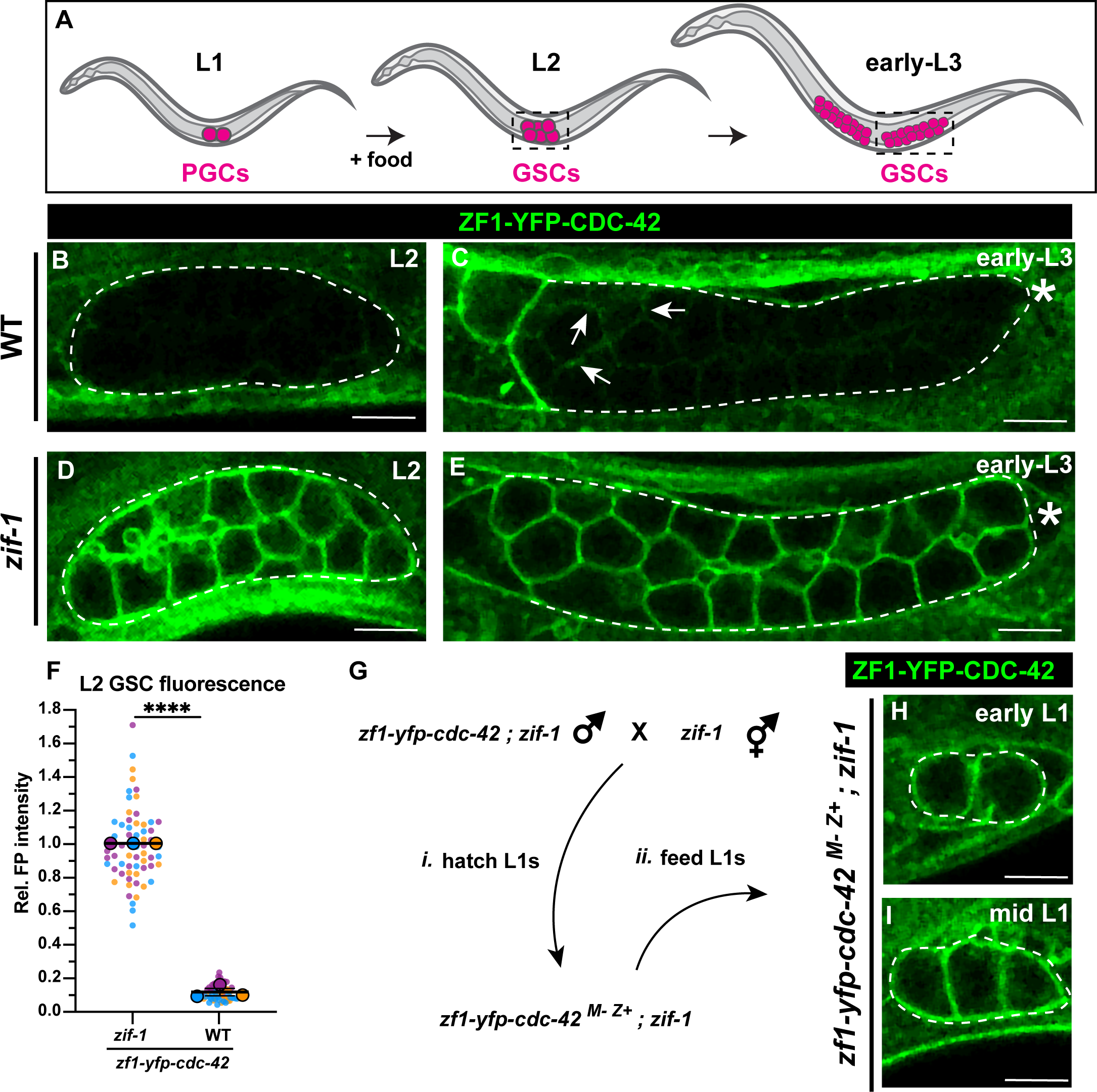
ZIF-1 mediates degradation in proliferative germ cells during early larval development. (**A**) Schematic of early larval germline development: Upon feeding, PGCs (magenta) re-enter the cell cycle and become undifferentiated germline stem cells GSCs (also magenta). GSCs continue to proliferate as development progresses. L2 and early L3 stages shown, dashed boxes represent the region of the worm imaged. (**B**) ZF1-YFP-CDC-42 is absent from wild-type L2 larval GSCs (dashed lines). (**C**) Expression of ZF1-YFP-CDC-42 in wild-type is first detected in early L3 GSCs, but is restricted to the proximal gonad (arrows), ZF1-YFP-CDC-42 signal remains weak near the distal gonad (asterisks). (**D-E**) ZF1-YFP-CDC-42 is expressed throughout the gonad in *zif-*1 mutant L2 and early L3 GSCs. (**F**) Quantification of L2 GSC ZF1-YFP-CDC-42 fluorescence in WT and *zif-1* mutants, normalized to expression in *zif-1* mutants. **(G)** Crossing scheme to introduce ZF1-YFP-CDC-42 zygotically in PGCs. (**H-I**) ZF1-YFP-CDC-42 expression in fed L1 larval PGCs. Individual data points from three independent color-coded biological replicates are shown as small dots, the mean from each experiment shown as a larger circle, the mean of means as a horizonal line, and the S.E.M as error bars. \*\*\*\**p* ≤ 0.0001, unpaired two-tailed Student’s *t*-test. Scale bar, 5µm.

We hypothesized that prior to or upon the PGC-to-GSC transition, ZIF-1 would become inactive and ZF1-tagged proteins would reappear in the germ line as a result of their zygotic expression. To test this hypothesis, we observed the expression pattern of ZF1-YFP-CDC-42 in fed worms at different larval stages. Contrary to our expectations, ZF1-YFP-CDC-42 remained nearly undetectable in wild-type gonads through the L2 stage (**FIG3B**). By the early L3 stage, we observed some re-expression of ZF1-YFP-CDC-42, but the protein was limited to the more proximal region of the gonad (**FIG3C, arrowheads**), corresponding approximately to where GSCs first enter meiotic prophase (Pepper *et al*. 2003; Hansen *et al*. 2004). This pattern of ZF1-YFP-CDC-42 absence mirrors the expression of *pie-1*p::*GFP-H2B*::*zif-1^3’UTR^*in the gonad, which turns on when L1 PGCs reenter the cell cycle and becomes restricted to undifferentiated germ cells by the early L3 larval stage (**FIGS1**) (Guven-Ozkan *et al*. 2010; Roy *et al*. 2018). By contrast, ZF1-YFP-CDC-42 was robustly expressed throughout L2 and early-L3 larval gonads in *zif-1* mutant worms (**FIG3D-F**). To exclude the possibility that ZF1-YFP-CDC-42 is absent in early larval GSCs because it is not yet transcribed, we crossed *zif-1* mutant mothers with *zif-1* mutant males containing the *zf1-yfp-cdc-42* allele to generate *zf1-yfp-cdc-42(M-Z+)* GSCs that lack endogenous *zif-1* activity (**FIG3G**). ZF1-YFP-CDC-42 rapidly appeared in fed *zif-1* mutant, *zf1-yfp-cdc-42(M-Z+)* gonads by the early L1-L2 stage (**FIG3H-I**) (20/21 early L1; 10/10 mid-L1 to L2), confirming that ZF1-YFP-CDC-42 is zygotically expressed in GSCs. These findings indicate that ZIF-1 in distal regions of the larval gonad clears ZF1-YFP-CDC-42 protein from undifferentiated germ cells.

The *pie-1*p::*GFP-H2B*::*zif-1^3’UTR^*transgene continues to be expressed in undifferentiated germ cells in the adult germline (**FIGS1**) (Guven-Ozkan *et al*. 2010; Roy *et al*. 2018). raising the possibility that ZIF-1 remains constitutively active in undifferentiated germ cells. To investigate ZIF-1 activity in adults, (**FIG4A**), we examined three different ZF1-tagged proteins: endogenous ZF1-YFP-CDC-42 and ZF1-YFP-CED-10, and transgenic *pie-1*-driven mCherry-PH-ZF1. Levels of each ZF1-tagged protein in the distal-most germ cells were severely depleted in a *zif-1*-dependent manner, though less completely than in PGCs or L2 larval undifferentiated germ cells (**FIG4B-G**). Taken together, these findings indicate that ZIF-1 switches roles from clearing proteins containing the ZF1 domain from early embryonic somatic cells, to clearing the same proteins from undifferentiated germ cells throughout the development of the germ line. This interpretation is consistent with the described spatial expression patterns of endogenous CCCH Zinc finger domain germ line proteins that are known targets of ZIF-1 (PIE-1, MEX-1, MEX-5, and POS-1), which are detected in the proximal but not distal regions of the gonad (Guedes and Priess 1997; Tenlen *et al*. 2008; Tsukamoto *et al*. 2017; Kim *et al*. 2021). We postulate that ZIF-1 could function redundantly with 3’ UTR mediated-translational regulation (Merritt *et al*. 2008) to ensure these spatial expression patterns within the germline syncytium (<u>see</u> <u>Discussion</u>).

**Figure 4.**
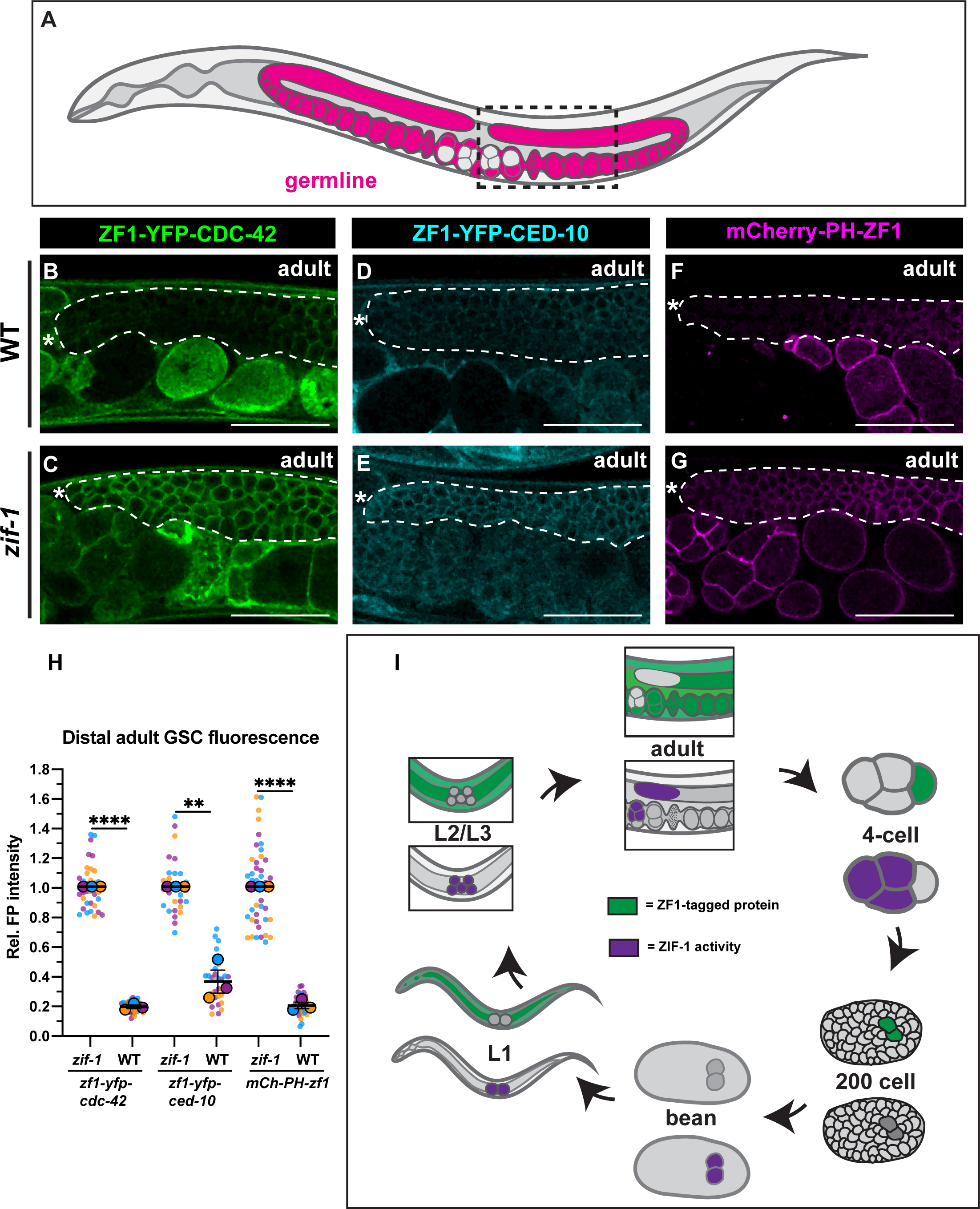
ZIF-1 remains active in proliferative germ cells through adulthood. (**A**) Schematic of the adult *C. elegans* germline. The dashed box represents the approximate region of the gonad that was imaged. (**B**) Localization pattern of ZF1-YFP-CDC-42 in wild-type adult GSCs. (**C**) Localization pattern of ZF1-YFP-CDC-42 in *zif-1* mutant adult GSCs. (**D**) Localization pattern of ZF1-YFP-CED-10 in wild-type adult GSCs. (**E**) Localization pattern of ZF1-YFP-CED-10 in *zif-1* mutant adult GSCs. (**F**) Localization pattern of mCherry-PH-ZF1 in wild-type adult GSCs. (**G**) Localization pattern of mCherry-PH-ZF1 in *zif-1* mutant adult GSCs. **(H)** Quantification of fluorescence in adult GSCs in wild-type and *zif-1* mutants, normalized to expression in *zif-1* mutants. (**I**) Schematic summarizing ZIF-1 activity throughout the life-cycle of *C. elegans*. Individual data points from three independent color-coded biological replicates are shown as small dots, the mean from each experiment shown as a larger circle, the mean of means as a horizonal line, and the S.E.M as error bars. \*\**p* ≤ 0.01, \*\*\*\**p* ≤ 0.0001, unpaired two-tailed Student’s *t*-test. Scale bar, 50µm.

### Degradation of ZF1-tagged proteins in PGCs can produce loss-of-function phenotypes

Based on these findings, we wondered whether ZIF-1-mediated depletion of proteins tagged with the ZF1 domain could provide a means to generate mutant phenotypes in undifferentiated germ cells. To test this idea, we tagged the endogenous *nop-1* gene with sequences encoding the ZF1 domain. NOP-1 is a myosin regulator that was originally identified for its requirement to form a plasma membrane invagination called the pseudocleavage furrow in pronuclear stage embryos (Rose *et al*. 1995; Tse *et al*. 2012; Zhang *et al*. 2018). We previously showed that NOP-1 is also required to stabilize the incomplete cytokinetic furrow that separates the two PGCs (Maniscalco *et al*. 2020); as a result, most PGCs become binucleate by the L1 stage (**FIG5A-B, E**) (Schwartz *et al*. 2022). *zf1-nop-1* embryos successfully formed a pseudocleaveage furrow (20/20 embryos), indicating that ZF1-NOP-1 is functional in early embryos. However, *zf1-nop-1* L1 larvae displayed an identical binucleate PGC phenotype with comparable penetrance (98%) to *nop-1* null mutant animals (93%) (**FIG5C-D, E**). The binucleate phenotype was suppressed when *zf1-nop-1* was introduced into a *zif-1* mutant background (1% binucleate) (**FIG5D-E**), indicating that the binucleate phenotype results from ZF1-NOP-1 degradation. Thus, endogenous ZIF-1 activity in undifferentiated germ cells can be harnessed as a genetic tool to generate loss-of-function phenotypes.

**Figure 5.**
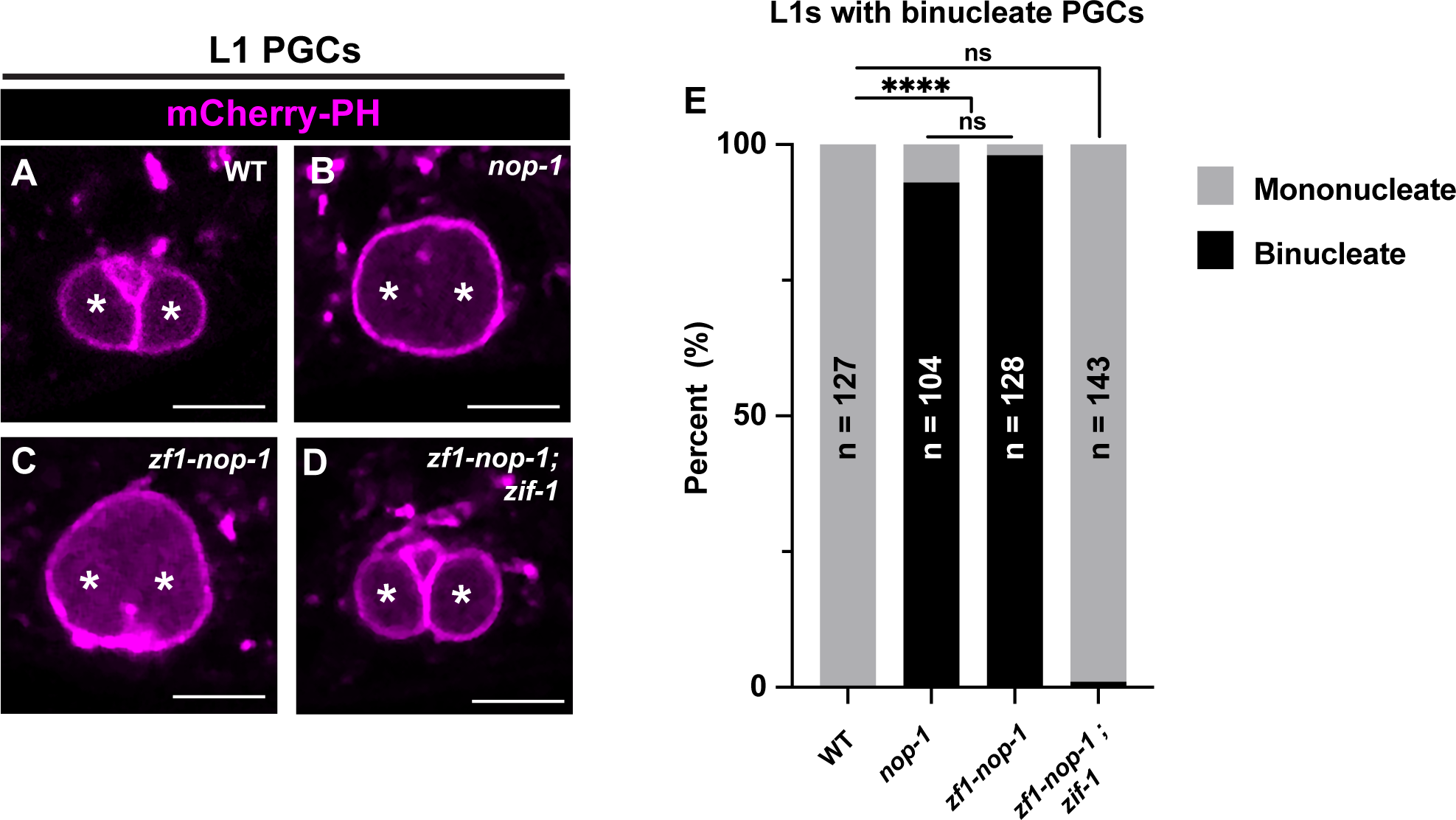
ZIF-1-mediated protein depletion is sufficient to generate PGC phenotypes. (**A-D**) Characterization of L1 larval PGC membrane morphology in wild-type, *nop-1*, *zf1-nop-1*, and *zf1-nop-1; zif-1* mutants. PGC nuclei denoted by asterisks. (**E**) Quantification of the binucleate PGC phenotype in wild-type, *nop-1*, *zf1-nop-1*, and *zf1-nop-1; zif-1* mutants. Fisher’s exact test was used to determine statistical significance. n.s., not significant (*p*> 0.05), **** *p* ≤ 0.0001. Scale bar, 5µm.

## Discussion

Our findings highlight a developmental role for *zif-1* in the *C. elegans* germ line (**FIG4I**). ZIF-1 was identified for its requirement to clear PIE-1 and several other CCCH Zinc finger proteins from early embryonic somatic cells, contributing to the enrichment of these proteins in germline precursor cells (Reese *et al*. 2000; DeRenzo *et al*. 2003). A maternal reporter transgene containing the *zif-1* 3’ UTR mirrors this soma-germline pattern of activity in its expression (Guven-Ozkan *et al*. 2010), suggesting that spatial differences in ZIF-1-mediated degradation in early embryos result from 3’ UTR-mediated regulation of maternal *zif-1* mRNA. Although PIE-1 normally disappears from PGCs soon after they are born, a previous study revealed that it perdured in PGCs in a *zif-1* mutant, indicating that *zif-1* has at least an indirect role in germline PIE-1 stability (Checchi and Kelly 2006). Here, by showing that heterologous proteins containing the PIE-1 ZF1 domain are cleared from PGCs in a *zif-1*-dependent fashion, we provide evidence that ZIF-1 becomes expressed in PGCs, at least in part through its zygotic transcription, and clears proteins containing the ZF1 domain. *zif-1* continues to be expressed in the germline of larvae and adults, where its activity is restricted to undifferentiated germ cells. This spatial control within the gonad is likely mediated by 3’ UTR translational regulation, since the *zif-1* 3’ UTR reporter shows an expression pattern in the gonad that mirrors where we show ZIF-1 to be active (Guven-Ozkan *et al*. 2010; Roy *et al*. 2018) (**FIGS1**). Thus, in addition to its previously described role in clearing PIE-1 and other CCCH Zinc finger proteins from early embryonic somatic cells, ZIF-1 subsequently removes these proteins from PGCs and undifferentiated germ cells (**FIG4I**).

We show that endogenous ZIF-1 activity can be harnessed to generate loss-of-function phenotypes in undifferentiated germ cells. Importantly, the ZIF-1/ZF1 degron system provides a method for cell-type specific genetic analysis in PGCs, which has not previously been possible. For example, since *zif-1* is transcribed zygotically in PGCs, one could study the function of an essential gene in PGCs by tagging it with sequences encoding the ZF1 domain within a *zif-1* mutant background; by crossing in a *zif-1(+)* allele from the male, ZIF-1 would become expressed for the first time in PGCs of F1 embryos and subsequently degrade the ZF1-tagged protein. This approach should similarly allow knockdown of proteins in undifferentiated germ cells of larvae and adults. As a corollary, however, geneticists using conditional ZIF-1 expression to degrade proteins in somatic cells should be aware that endogenous ZIF-1 will also target the protein in PGCs and undifferentiated germ cells. This caveat, which must be taken into account when interpreting phenotypes, could be circumvented by performing experiments in a *zif-1* mutant background, as has been employed when studying the role of essential genes during early embryonic development (Sallee *et al*. 2018).

Why is *zif-1* expressed in PGCs at all? One possibility could be to aid in the initiation of zygotic transcription in the early germ line. Though largely transcriptionally quiescent (Mello *et al*. 1996; Seydoux 1996; Seydoux and Dunn 1997), PGCs initiate the transcription of a small set of genes during mid-embryogenesis (Subramaniam and Seydoux 1999; Kawasaki *et al*. 2004; Mainpal *et al*. 2015; Belew *et al*. 2021). PGC zygotic genome activation coincides with the removal of PIE-1 from PGCs by ZIF-1 and is correlated with the reappearance of phosphorylated RNA pol II in PGCs (Seydoux and Dunn 1997); as suggested previously, ZIF-1 may function to enhance transcription in PGCs by removing PIE-1, a global RNAPII inhibitor (Checchi and Kelly 2006). However, PIE-1 should in theory also prevent zygotic *zif-1* transcription, which we show occurs in PGCs. One possibility is that maternal *zif-1* mRNA could be translated in young PGCs, which would initiate PIE-1 disappearance and allow zygotic expression of *zif-1* and other genes.

How maternal *zif-1* mRNA could switch from being translationally repressed in P1-P4, to being activated in PGCs is unclear. However, the *zif-1* translational repressor SPN-4 disappears from PGCs shortly after PGC birth via an unknown mechanism (Ogura *et al*. 2003; Oldenbroek *et al*. 2012). SPN-4 works in concert with POS-1 to repress *zif-*1 (Oldenbroek *et al*. 2012). We postulate that POS-1 alone may be insufficient to completely repress *zif-1* translation in PGCs, and SPN-4 disappearance could thus allow for translation of maternal *zif-1* mRNAs present in early PGCs. Consistent with this possibility, we observed modest translation of *pie-1*p::*GFP-H2B*::*zif-1^3’UTR^* in some early PGCs, indicating that the *zif-1* 3’ UTR is competent to promote some translation of maternal RNAs at this stage.

Although ZIF-1 is required for the timely removal of PIE-1 from PGCs, PIE-1 protein eventually dissipates regardless of the presence of functional ZIF-1 (Schaner *et al*. 2003; Checchi and Kelly 2006), suggesting either that ZIF-1 functions redundantly with an additional mechanism for PIE-1 removal, or that PIE-1 has a modest half-life and disappears through decay. In the early embryo, ZIF-1 clears a suite of CCCH Zinc finger proteins from somatic cells (Mello *et al*. 1996; Seydoux 1996; Guedes and Priess 1997; Tabara *et al*. 1999; DeRenzo *et al*. 2003), and we consider it likely that ZIF-1 clears these proteins from PGCs as well. Consistent with this idea, CCCH Zinc finger proteins POS-1 and MEX-1 are highly enriched in germline precursor cells but not in PGCs of late embryos (Guedes and Priess 1997; Tabara *et al*. 1999).

Accomplishing a clean transition from maternal-to-zygotic gene expression (MZT) is critical for proper embryonic development, and multiple mechanisms exist to achieve such a transition in both somatic and germline lineages (Vastenhouw *et al*. 2019). One recently described mechanism that regulates MZT in PGCs requires the E3 ligase component GRIF-1. Similar to ZIF-1, GRIF-1 becomes active in PGCs shortly after their birth, where it targets the maternally deposited cytoplasmic poly(A)polymerase complex catalytic subunit GLD-2 for degradation (Oyewale and Eckmann 2022). Loss of *grif-1* function results in an aberrant PGC mRNA expression profile and severe transgenerational defects (Oyewale and Eckmann 2022). PIE-1 disappearance from PGCs is unaffected in *grif-1* mutants (Oyewale and Eckmann 2022), suggesting that ZIF-1 and GRIF-1 act independently. We postulate that ZIF-1 and GRIF-1 function in parallel to optimize PGC MZT via the concurrent degradation of PIE-1 and GLD-2.

It is unclear why ZIF-1 continues to remain active in undifferentiated larval and adult germ cells. One possibility is that ZIF-1 could function redundantly with existing mechanisms that regulate the spatial distribution of germline proteins in the adult gonad. One such mechanism occurs at the translational level, where a myriad of RNA-binding proteins restricts mRNA translation to certain germline domains via binding of the 3’ UTR of transcribed mRNAs. Multiple CCCH Zn finger domain containing proteins, including PIE-1, are expressed in the adult germline, but are restricted to the proximal gonad (Guedes and Priess 1997; Detwiler *et al*. 2001; Kritikou *et al*. 2006; Tenlen *et al*. 2008; Tsukamoto *et al*. 2017; Kim *et al*. 2021), at least in part because of 3’ UTR regulation (Merritt *et al*. 2008). However, ZIF-1 could function as a redundant mechanism to ensure that PIE-1 and other direct ZIF-1 targets (MEX-1, POS-1, and MEX-5) are absent in undifferentiated germ cells (Guedes and Priess 1997; Tsukamoto *et al*. 2017; Kim *et al*. 2021). Such a role may have evolved as a result of the syncytial organization of the germ line, where germ cells are connected to a common cytoplasmic core. At least in principle, proteins expressed in one region of the gonad could diffuse to others. If this were to occur with other CCCH Zinc finger proteins, ZIF-1 could ensure that, in combination with 3’ UTR regulation, they were efficiently excluded from undifferentiated germ cells.

## Data availability

All strains, plasmids and sequences are available upon request. All data supporting the conclusions in the manuscript are included within the text, figures, and tables.

## Acknowledgements

We thank the *Caenorhabditis* Genetics Center (CGC), Ann Wehman (University of Denver), Bob Goldstein (UNC), and E. Jane Hubbard (NYU School of Medicine) for strains. The CGC is supported by the NIH Office of Research Infrastructure Programs (P40OD010440). We thank the members of the Nance laboratory for comments on the initial manuscript. Microscopy used instrumentation in the NYULMC Microscopy Laboratory, which is partially supported by the Laura and Isaac Perlmutter Cancer Center support grant P30CA016087 from the National Institutes of Health/National Cancer Institute. This work was supported by a training grant from NYSTEM (C32560GG) and a fellowship from the National Institutes of Health (NIH) (F31HD102161) to AZAS; and a research grant from the NIH to JN (R35GM118081).

## Author contributions

AZAS and JN conceived the project. AZAS performed all experiments except creation of the *zf1-yfp-ced-10* allele, which was made by YA. AZAS and JN analyzed and interpreted the data. AZAS and JN wrote the initial manuscript, and all authors edited the manuscript.

## Methods

### Worm culture and strains

All strains were maintained at 20°C and raised on NGM plates seeded with *Escherichia coli* strain OP50 (Brenner 1974). For larval imaging experiments, synchronized L1 larvae were isolated by treating gravid adults with alkaline hypochlorite and hatching eggs in M9 buffer overnight. Synchronized L1s were then fed either 24, 36 and 72 hours at 20°C on enriched peptone plates seeded with *E. coli* strain NA22 to isolate L2 larvae, L3 larvae and adults, respectively. A list of all strains used in the study can be found in Supplementary Table 1.

### Microscopy

Embryos, adults, and larvae were mounted on 5% agarose pads. Larvae and adults were immobilized prior to and during image acquisition using 1.25mM levamisole in M9 buffer. Animals were imaged on a Leica TCS SP8 laser-scanning confocal microscope using a 63X 1.4 NA oil-immersion objective with 488nm and 594nm lasers and HyD detectors. Images were analyzed and processed in ImageJ (NIH) and Photoshop (Adobe).

### Image analysis

ImageJ was used to measure the fluorescence intensity of ZF1-tagged proteins. For PGCs of bean stage embryos and starved L1 larvae, measurements were taken from a single 6-pixel line drawn along the contact site of the two PGCs. For L2 larvae, measurements were taken from a region of interest encompassing the whole L2 germ line. For adults, measurements were taken from a region of interest encompassing ∼10 cell diameters from distal-most adult germ line. No background subtractions were made before data analysis.

### Zygotic gene expression experiments

To determine if zif-1 was expressed zygotically in PGCs, hmr-1-mKate males were crossed to zf1-yfp-cdc-42; zif-1(gk117) mutant hermaphrodites. Cross-progeny, which were distinguished from self-progeny by somatic zygotic expression of HMR-1-mKate in late-stage embryos and L1 larvae, were scored for detectable expression of ZF1-YFP-CDC-42 in the PGCs.

To determine when zf1-yfp-cdc-42 zygotic expression began in the larval germ line, zf1-yfp-cdc-42; zif-1(gk117) males were crossed to zif-1(gk117) hermaphrodites. F1 progeny were allowed to feed on the cross plate for 6-12 hours before they were washed off and mounted on slides as described above. Cross-progeny, which were identified by somatic expression of ZF1-YFP-CDC-42, were scored for detectable expression of ZF1-YFP-CDC-42 in L1 germ cells.

### Generation of endogenously tagged alleles

#### zf1-ha-nop-1

Pre-incubated Cas9 (Berkeley) and crRNA + tracrRNA (IDT) ribonucleoprotein was combined with a ssDNA oligo (IDT) (with ∼25-35bp homology arms) repair template to generate edits. Candidate edited worms were identified using the dpy-10(cn64) co-CRISPR strategy, as previously described (Arribere et al. 2014; Paix et al. 2017). Two independent zf1-ha-nop-1 alleles were isolated. To generate zf1-ha-nop-1 in a wild-type background, nop-1(xn194[zf1-ha-nop-1]) was generated by directly injecting into FT1900, which contains a PGC-specific membrane marker. To generate zf1-ha-nop-1 in the zif-1(gk117) background, an additional allele, nop-1(xn200[zf1-ha-nop-1]) was generated via direct injection into zif-1(gk117) worms.

#### zf1-yfp-ced-10

Cas9 and a sgRNA were expressed from a single plasmid by replacing the sgRNA homology sequence in pDD122 (Dickinson 2013) with an sgRNA sequence targeting the 5’ end of ced-10. zf1-yfp was introduced from a plasmid containing 500bp homology arms to create zf1-yfp-ced-10. The unc-119(+) gene was included as a co-transformation marker in reverse orientation within a yfp intron, and DNAs were injected into unc-119(ed3) mutant worms as described (Armenti et al. 2014; Zilberman et al. 2017). Non-Unc progeny were screened by fluorescence and PCR to identify the ced-10(xn53[zf1-yfp-ced-10) allele.

### Statistical analysis

Statistical analysis was performed using GraphPad Prism 9 software. For categorical data, such as scoring the binucleate phenotype in nop-1 L1 larval PGCs, contingency tables were generated and Fisher’s exact test was performed. For all other data, unpaired two-tailed students t-tests were performed. The data in graphs are represented as Superplots (Lord et al. 2020), with individual data points from three independent color-coded biological replicates shown as small transparent dots, the mean from each individual experiment shown as larger circles, the mean of means shown as a horizontal line, and error bars representing the standard error of the mean (S.E.M). Statistical test type, sample size, and p-values are reported in figure legends.

---

**Figure S1.**
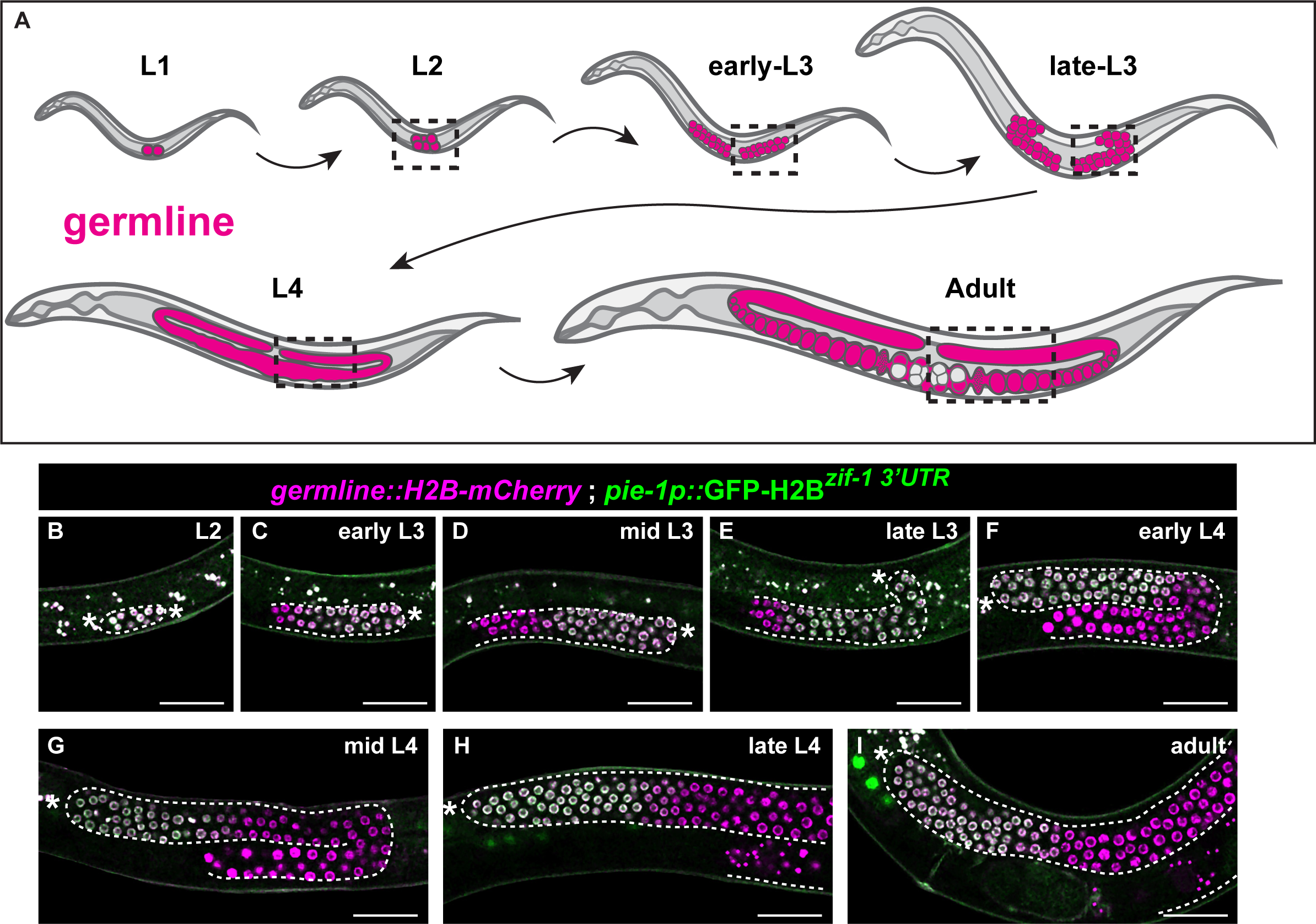
A *zif-1 3’UTR* reporter is restricted to proliferative germ cells at all stages of germline development. (**A**) Schematic of germline proliferation from L1 PGCs to the entire adult germline. Two gonad arms proliferate in opposite directions and by late L3 turn back towards the worm midline. Proliferative germ cells are restricted to the distal gonad as it elongates in each gonad arm. Dashed boxes represent the region of the worm imaged. (**B-I**) *pie-1p::H2B-GFP^zif-1^ ^3’UTR^* is expressed in L2 larval GSCs, after which it becomes restricted to undifferentiated (distal) germ cells and remains expressed in these cells through adulthood. All germ cells marked with *germline*::*mCherry-H2B* (*mex5p::mCherry-H2B^nos-2^ ^3’UTR^*). Distal end of the gonad denoted by asterisks. Scale bar, 25 µm

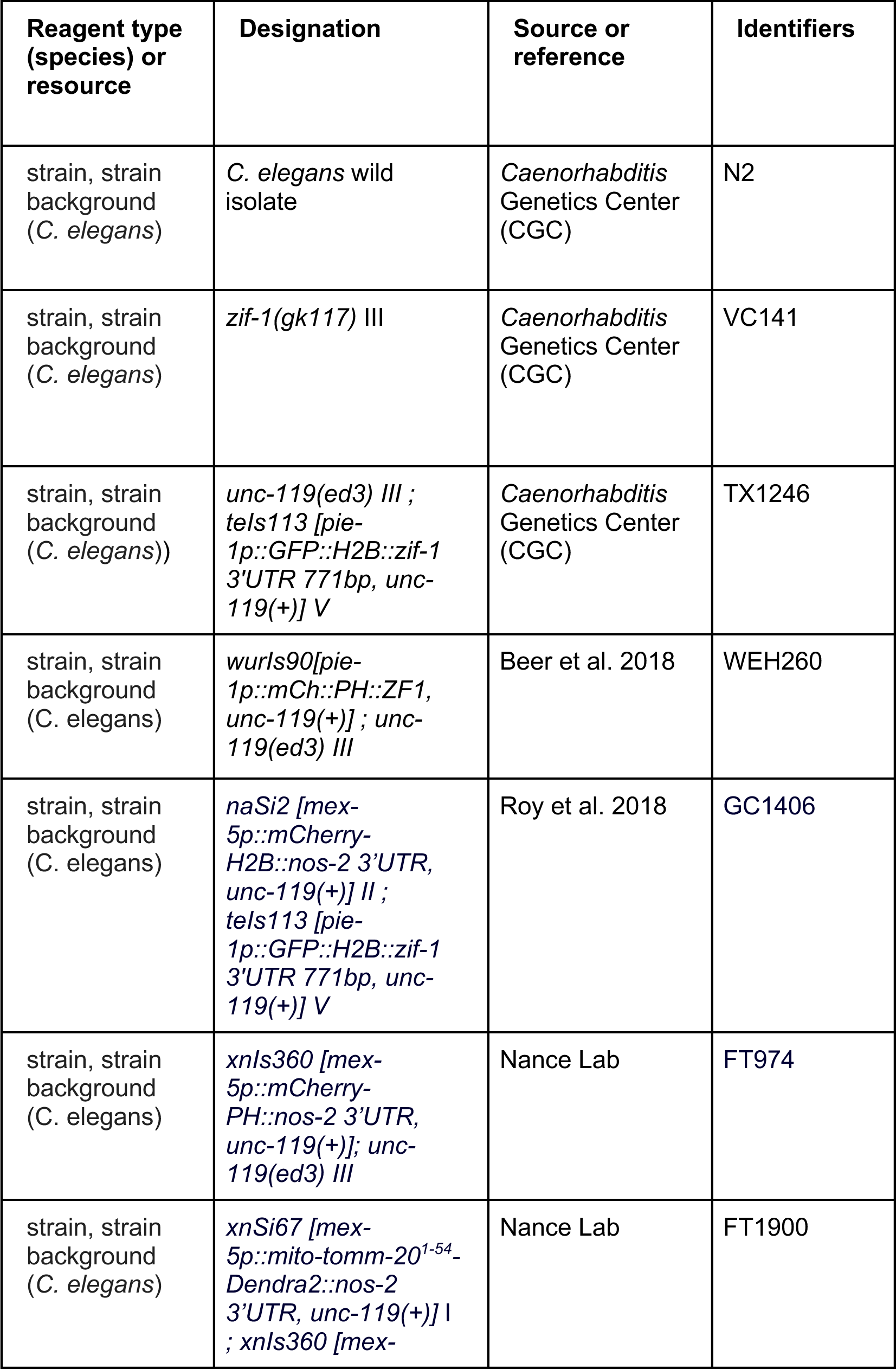
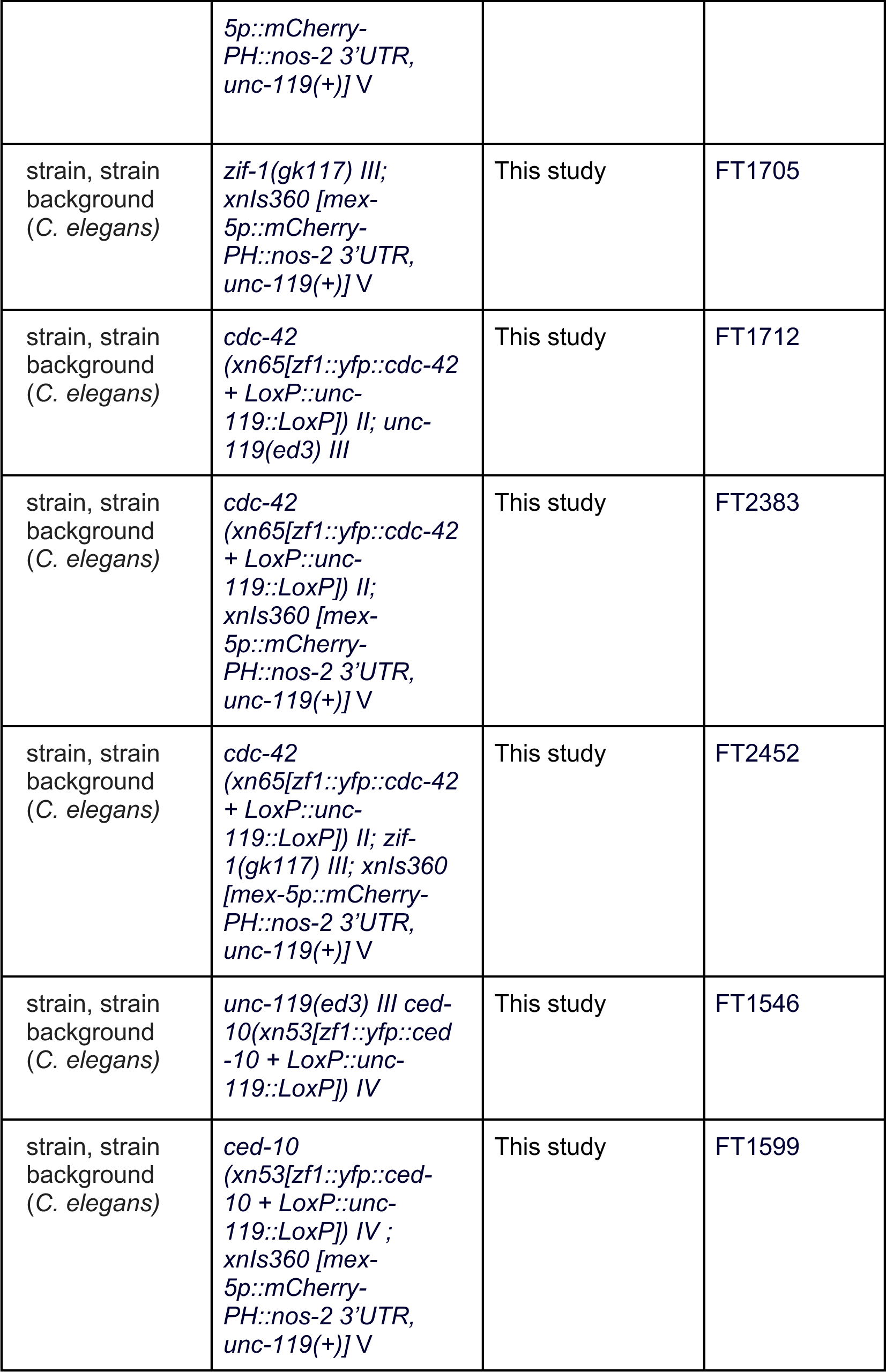
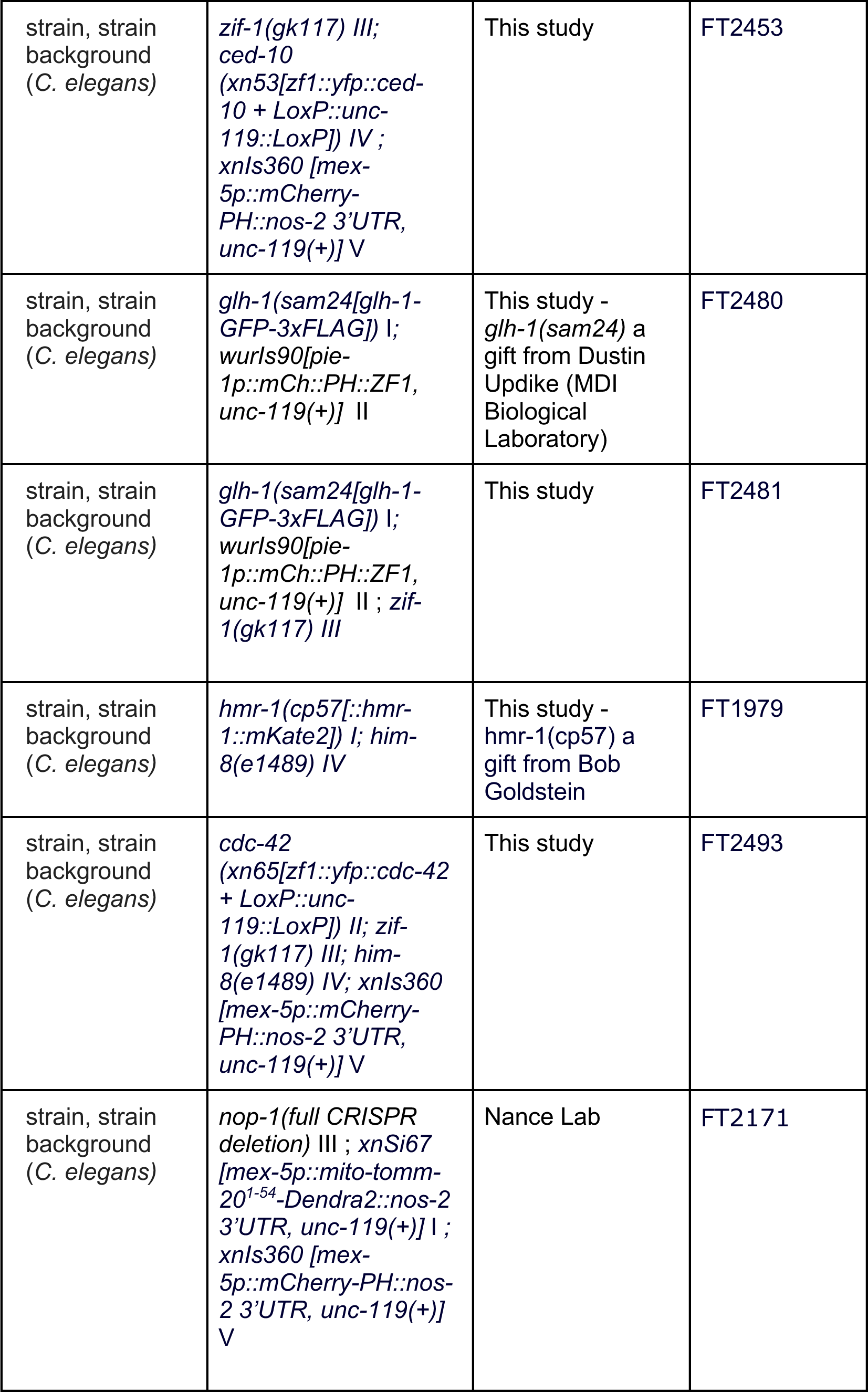
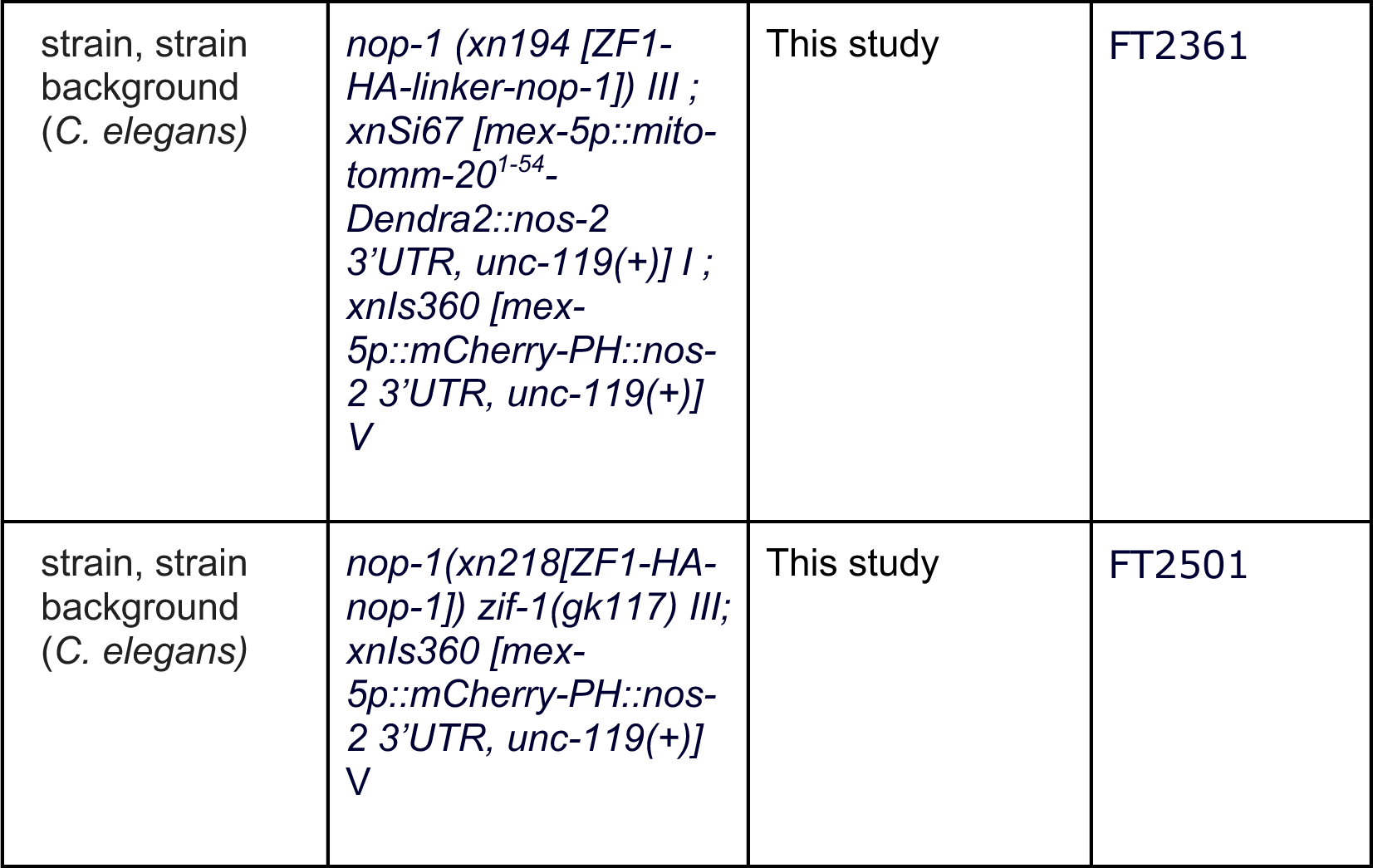
Supplemental Table 1

